# Isolation of the side population from adult neurogenic niches enriches for endothelial cells

**DOI:** 10.1101/2021.04.16.440120

**Authors:** Alena Kalinina, Catherine Gnyra, Yingben Xue, Diane Lagace

## Abstract

In stem cell research, DNA-binding dyes offer the ability to purify live stem cells using flow cytometry as they form a low-fluorescence side population due to the activity of ABC transporters. Adult neural stem cells exist within the lateral ventricle and dentate gyrus of the adult brain yet the ability of DNA-binding dyes to identify these adult stem cells as side populations remain untested. The following experiments utilize the efflux of a DNA-binding dye, Vyrbant DyeCycle Violet (DCV), to isolate *bona fide* side populations in the adult mouse dentate gyrus and SVZ and test their sensitivity to ABC transporter inhibitors. A distinct side population was found in both the adult lateral ventricle and dentate gyrus using DCV fluorescence and forward scatter instead of the conventional dual fluorescence approach. These side populations responded strongly to inhibition with the ABC transporter antagonists, verapamil and fumitremorgin C. The cells in the side population were identified as cerebrovascular endothelial cells characterized by their expression of CD31. These findings, therefore, suggest that the side population analysis provides an efficient method to purify endothelial cells, but not adult neural stem cells.

## Introduction

DNA-binding dyes have been perpetually used in flow cytometry and fluorescence-activated cell-sorting (FACS) paradigms to identify cancer stem cells (1,2). This has included the use of dyes such as Hoechst 33342 (3,4) and, more recently, Vybrant DyeCycleViolet (DCV), which is less toxic to stem cells (8–10). In these assays, live stem cells are identified as a side population that has low dual fluorescence intensity in both blue- and red-shifted parameters due to the activity of ABC transporters, which can efflux the DNA-binding dyes. In contrast, cells that do not have ABC transporters will accumulate the dye and show higher fluorescence. Since ABC transporters have been identified in stem cells from a large variety of tissues (8), this method has extended to be used routinely to isolate various types of stem cells.

The isolation of neural stem cells (NSCs) in the subventricular zone (SVZ) of the lateral ventricle and the subgranular zone of the dentate gyrus has been a challenge in the field due to many reasons. These include the relatively small number of NSCs, the multiple subtypes of NSCs, and lack of a highly specific surface marker for identification and isolation by FACS (9,10). Classically, flow cytometry paradigms utilized low expression of PNA (peanut agglutinin) and HAS (heat stable antigen), or high expression of Notch, LewisX (cluster of differentiation-15, CD15) and EGFR (early growth response factor) surface markers to identify neural stem cells (11–14). More recent methods have made additional significant advances in identification and enrichment of subpopulations of NSCs using multi-parameter FACS, inducible transgenic mice models (12), or single-cell transcriptional analyses (15–18). However, these methodologies are time- and cost-intensive, which has led our lab and others to investigate the use of DNA-binding dyes as a simpler and more efficient method for identification and purification of NSCs

Many have identified an ABC transporter-dependent side population with an NSC identity in cells isolated from neurospheres derived from primary embryonic neural or postnatal/adult SVZ tissue (3,11,19). In contrast, NSCs isolated *ex vivo* in cells freshly harvested from embryonic or early postnatal SVZ (postnatal day 2) brain tissue are not found in the side population (3,11,19). Instead of NSCs, endothelial and microglial cells were comprising the side population identified in *ex vivo* preparations of developing SVZ (3). This raises the question of whether NSC-containing side populations can be identified from *ex vivo* primary adult mouse dentate gyrus and SVZ tissue. To answer this question, we optimized the detection and phenotyping of the side population using flow cytometry and the DNA-binding dye, DCV, in live single-cell suspensions from the adult mouse dentate gyrus and SVZ. The data shows that a ABCG2/B1-dependent side population can be identified in the adult neurogenic niches, that is enriched for endothelial cells but not NSCs.

## Materials and Methods

### 1. Animals

This study was carried out in strict accordance with the recommendations in the Guidelines of the Canadian Council on Animal Care and all efforts were made to minimize suffering. The animal care protocol was approval by the University of Ottawa Animal Care Committee (Protocol CMM-1150). Fifty two- to three-month-old male and female mice (C57bl/6J background) were used for all experiments. Animals were group housed in standard laboratory cages and kept on a 12-hour night/day cycle with *ad libitum* access to food and water.

### 2. Tissue collection and digestion

Mice were deeply anesthetized with euthanyl (90 mg/kg) and the brains were quickly placed in ice-cold artificial cerebrospinal fluid (aCSF, pH = 7.4) prepared in miliQ water with 124mM NaCl, 5mM KCl, 1.3 mM MgCl2·6H2O, 2mM CaCl2·2H2O, 26mM NaHCO3, and 1X penicillin-streptomycin (10,000 U/mL; ThermoFisher) and sterilized using stericup and steritop filtration set (Millipore). Dentate gyrus and SVZ were microdissected using SteREO Discovery V8 microscope (Zeiss) following previously published protocols (20,21).

Tissue was digested according to protocols described previously (22,23). First, the tissue was gently broken up using small surgical scissors then incubated on twister (30 minutes, 37oC) in 500uL of digestion media, containing 20 U/mL papain (Worthington Biochemicals), 12 mM EDTA (Invitrogen) in DMEM:F12 (Invitrogen). Following incubation, Resuspension media (0.05 mg/mL DNase1 (Roche) with 10% fetal bovine serum (Wisent Bioproducts) in DMEM:F12) was added to each tube, triturated 10X with a P1000 micropipette, and incubated for five minutes at RT. Suspension was then transferred in Percoll media, consisting of 19.8% Percoll (GE Healthcare Life Sciences), 2.2% 10X PBS (Wisent Bioproducts) in Resuspension media. Cells were then spun down (500 x g, 13 minutes, 4oC), the supernatant was removed, and cells were resuspended in 200μL of phenol-free DMEM:F12. Live cells were counted on Countess automated cell counter (ThermoFisher Scientific) using 0.4% Trypan blue (Invitrogen) at a concentration of 1:2 and suspended in phenol-free DMEM:F12 medium at a concentration of 10^6^/mL.

### 3. Staining and drug treatments

Vybrant DyeCycle Violet Ready Flow™ Reagent (Invitrogen) was added to cells in phenol-free DMEM:F12 medium and incubated at 37oC in a 5% CO2 cell culture chamber (Forma Series II Water Jacket; ThermoFisher Scientific) for 30 minutes. The concentration of DCV was tested at both 1X and 2X, and based on these experiments (Fig S1), all future experiments used the concentration of 2X, or 160uL in 10^6^ cells/ml.

For experiments involving ABC transporter inhibition, fumitremorgin C (FTC; Sigma) and (±) verapamil hydrochloride (VP; Sigma) were added to unwashed cells at final concentrations of 10uM and 50uM, respectively, after DCV incubation and kept in the same conditions for additional 30 minutes. Cell were then kept on ice in dark until sort, and 7-Amino-Actinomycin D (7AAD, 40 ug/ml, Sigma) was added to cell suspensions 10 minutes before analysis for dead cell discrimination. For experiments determining the identity of the side population, CD31 antibody conjugated to allophycocyanin (APC), BD Biosciences, BioLegend), was added to cells in DMEM:F12 at final concentration of 1:50 (24) and incubated on ice in the dark for 30 minutes before DCV incubation, which followed the same workflow as discussed above. Antibodies, dyes, and drugs used for all experiments are listed in Table 1.

**Table 1.** Reagents used for tissue processing and DCV assay.

| Reagents | Company | Catalogue # | Final Concentration |
| --- | --- | --- | --- |
| Vybrant DyeCycle Violet Ready Flow™ Reagent | Invitrogen | R37172 | 160 uL/ml |
| APC anti-mouse CD31 | BioLegend | 102409 | 1:50 |
| 7AAD | Sigma | A9400-1MG | 1 ug/ml |
| Fumitremorgin C | Sigma | F9054-250UG | 10uM |
| (+/-) Verapamil Hydrochloride | Sigma | V4629-1G | 50uM |
| Papain suspension | Cedarlane | LS003126 | 20 U/ml |
| Percoll | Sigma | 17-0891-02 | 22% v/v |
| Trypan Blue | Invitrogen | T10282 | 1:1 (0.2%) |

### 4. Cell lines

Two cancer cell lines, U-2OS (ATCC, osteosarcoma) and A2780 S (17, ovarian cancer), were generously provided by Dr. Laura Trickle-Mulcahy and Dr. Barbara Vanderhyden, respectively. Cells were grown in DMEM/10%FBS until minimum 75% confluency was reached. Cells were detached from flasks in 5mM EDTA for 20 minutes at 37oC in a cell culture incubator, then triturated and washed several times with 1X PBS before cell count and DCV staining, which followed the same procedure as staining in primary brain cells.

### 5. Flow cytometry

All flow cytometry experiments were performed using BD LSRFortessaTM flow cytometer (BD Biosciences) in the Flow Cytometry and Virometry Core at the University of Ottawa, Faculty of Medicine. Unstained and single-stained controls were used to set up laser parameters and gating for all-stained samples. First, cell debris and doublets were excluded based on FSC and SSC parameters, and then 7AAD+ dead cells were removed from analyses. Following this, all samples were collected under 405nm laser with 450/50 and 660/20 bandpass filters. DCV+ populations could only be resolved with optimal excitation of the samples (Fig S1). 7AAD signal was collected under the 561nm laser with a 670/30 filter. APC-CD31 fluorescence was collected using the 640nm laser with a 660/20 bandpass filter without need for compensation. Single-stained controls were used to identify and gate CD31+ and CD31-cells. The side population fidelity of DCV+ cells was determined by comparison to FTC- and VP-treated samples.

### 6. Data analysis

FlowJo software (BD Biosciences) was used to analyze and visualize all flow cytometry data.

## Results

### Primary cells isolated from the dentate gyrus and SVZ contain multiple populations with side population properties

We used the Vybrant DyeCycle Violet Ready Flow Reagent™ (Invitrogen) to test the presence of a side population that was able to efflux the DNA-dye. Primary live cells harvested from the dissected neurogenic regions of the dentate gyrus and SVZ showed heterogeneous populations of DCV-stained cells as demonstrated by variable DNA content (Fig 1A and 1B). In both the dentate gyrus and SVZ populations there was a large population of cells with low DCV fluorescence that appeared in the lower left corner of dual fluorescence DCV-Blue/DCV-Red plots (Fig 1A and 1B). These cells in the lower corner resembled effluxing cells, which were absent in the negative control U2OS cell line (Fig 1C) that has been previously reported to not contain a side population (26,27).

**Fig 1.**
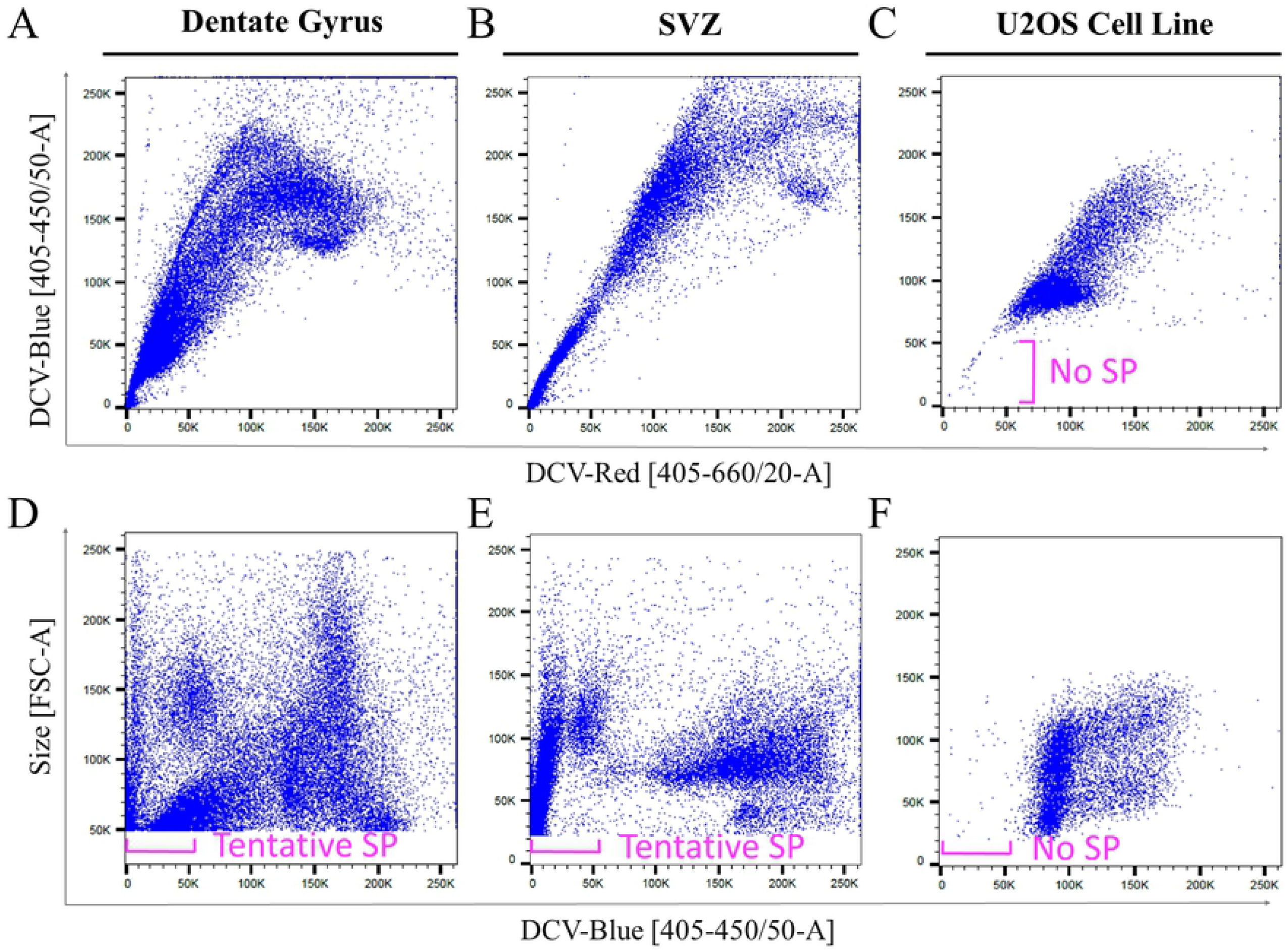
DCV identifies heterogeneous cell populations in primary neurogenic brain cells. Cell heterogeneity is illustrated in dual fluorescence DCV-Red vs DCV-Blue plots with adult cells isolated from the dentate gyrus (A), SVZ (B), as well as control cultured U2OS cells (C) that do not have a side population. Forward scatter (size) and DCV-Blue combined plots for dentate gyrus (D) and SVZ (E) show multiple low-fluorescence populations that could be bona fide side populations (labeled as Tentative SP), whereas, U2OS cells (F) show a few scattered cells that are debris and nuclear fragments.

In the cell suspensions from the dentate gyrus and SVZ it was difficult to distinguish the side population from the continuous main population using the red and blue dual fluorescence of DCV. Therefore, a forward scatter parameter (cell size) was added for the remainder of our analyses to observe the cell heterogeneity together with DCV fluorescence (Fig 1D-F). The size/DCV-Blue plots of dentate and SVZ cells reveal multiple low-fluorescence cell populations that appeared to be effluxing DCV, which we labeled as the tentative side population (Fig 1D and 1E). As expected the DCV-stained live U2OS control cells showed no low-fluorescence cell populations (Fig 1F), highlighting the lack of side population in this culture system.

### Side populations in primary dentate gyrus and SVZ are responsive to ABC transporter inhibition

To further identify the side population of the adult neural cells, fumitremorgin C (FTC) and verapamil (VP) were used to inhibit the activity of ABCG2 and ABCB1 transporters, which are known to prevent the efflux of DCV (6,28). Primary dentate gyrus cells located in the labeled tentative SP area in the size-DCV-blue plot, showed a population of cells that was reduced from 4.03% to 0.34% after addition of ABC-transporter antagonists (Fig 2A, 2D). Similarly, primary SVZ cells located in the tentative SP area responded to the inhibitor treatment with a reduction of the population from 0.71% to 0.22% after treatment (Fig 2B, 2E). To confirm specificity of ABC transporter inhibitors, the A2780 S cell line was used as a positive control for VP- and FTC-sensitive side population cells (2,7,29). As predicted, A2780 S cells had a very distinct side population in standard dual-fluorescence plots that was reduced from 7.97% to 1.51% after treatment (Fig 2C, 2F). Overall, these data suggest that primary dentate gyrus and SVZ cells contain a population of cells that possesses side population properties and effluxes DNA-binding dyes via ABC transporters, and that can be clearly visualized through measuring cell size together with the fluorescence of the DNA-binding dye.

**Fig 2.**
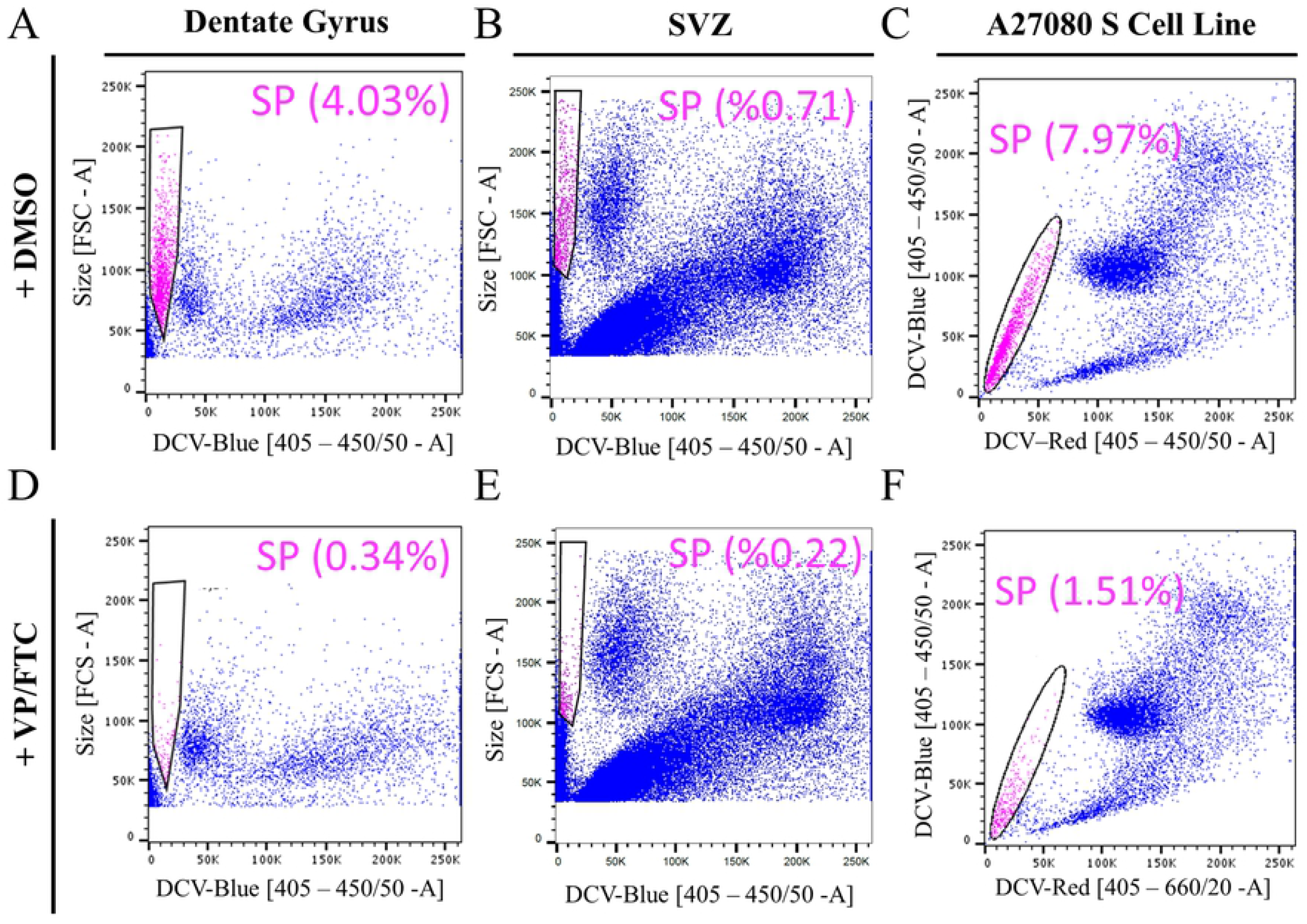
ABC transporter antagonists (VP and FTC) inhibit the side population phenotype of primary adult dentate gyrus and SVZ cells. Fluorescence/size plots for dentate gyrus (A) and SVZ (C) show the side population (SP, pink) and percentage of cells in the SP in the absence (A, B) and presence of VP and FTC (D,E) which significantly reduced the side population. A2780 S cell line, a positive control, shows a large side population in dual fluorescence plots (C) and nearly complete removal (F) of the effluxing side population by addition of inhibitors.

### The dentate gyrus and SVZ side populations comprise endothelial cells

We hypothesized that the side population may have endothelial cell identity, as have been previously identified in *ex vivo* early postnatal SVZ cells (3). This hypothesis was tested using the surface marker for endothelial cells, cluster of differentiation 31 (CD31) to identify them among the dentate gyrus cell types. The CD31 expressing (CD31+) endothelial cells represented 5.96% of all cells in the dissected adult dentate gyrus (Fig 3C) and 3.07% of all cells in the dissected adult SVZ (Fig 3F). All other cells, including the adult neural stem cells, were labeled as CD31-negative (CD31-) main population. CD31-cells from the dentate gyrus did not contain the majority of side population, as shown by the size-DCV-Blue plot (Fig 3A). On the contrary, CD31+ endothelial cells were heterogeneous in size and DNA content but mostly exhibited a homogenous low DCV fluorescence (Fig 3D). In fact, 75.48% of SP cells were CD31+ endothelial cells (Fig 3C). Similarly, for the SVZ, CD31-non-endothelial cells from the adult SVZ did not contain the majority of side population cells (Fig 3B), while CD31+ endothelial cells occupied this area (Fig 3E) and represented 52.65% of the side population (Fig 3F). This data, therefore, strongly suggests that the CD31+ endothelial cells mainly comprise the low-fluorescence DCV-effluxing cells that induce the side population phenotype of adult primary dentate gyrus and SVZ cells.

**Fig 3.**
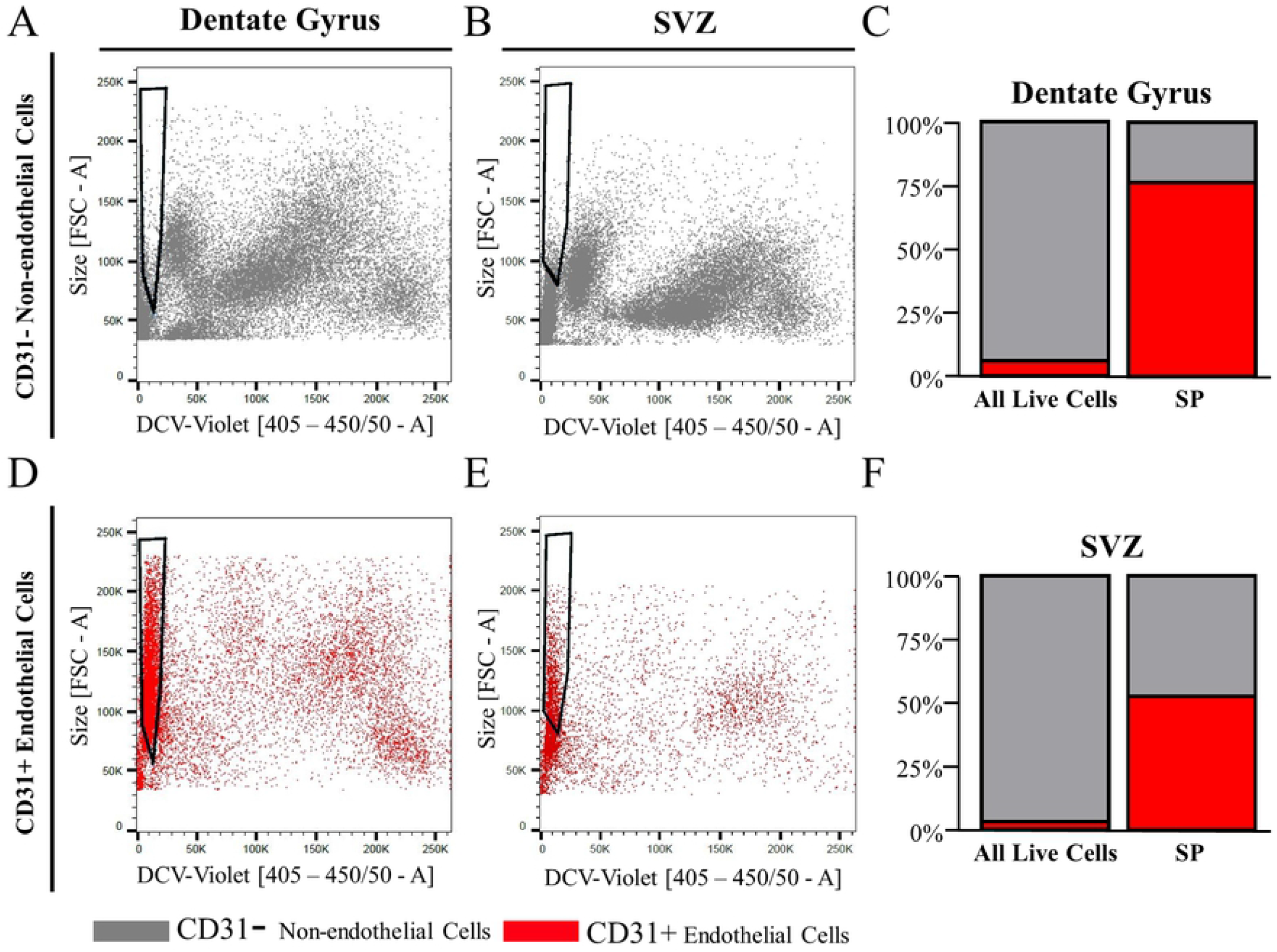
The side population phenotype of adult primary dentate gyrus and SVZ cells is mainly composed of endothelial cells. The CD31-negative cells are a heterogeneous population in the dentate gyrus (A) and SVZ (B) and do not contain the DNA-dye-effluxing side population. Alternatively, CD31-positive cells localize to the side population area make up a large portion of the SP in primary dentate gurus (D, C) and SVZ cells (E, F).

## Discussion

This series of experiments was used to examine whether a side population can be identified in primary cells of the dentate gyrus and SVZ using DCV, a live-cell-permeable DNA-binding dye, in flow cytometry. We identified a side population that effluxes DCV that was best visualized through measuring cell size together with the fluorescence of the DNA-binding dye. These cells represent the *bona fide* side population that responds to the inhibition with ABC transporter antagonists, verapamil and fumitremorgin C. In both the dentate gyrus and SVZ cells, the side population was largely composed of CD31-expressing cells, thus supporting the notion that side populations from adult neurogenic regions are of endothelial origin.

### Primary endothelial cells form the side population in ex vivo adult neurogenic extracts

Our data support that within the adult dentate gyrus and SVZ, endothelial cells are the major cell cluster in the side population that responded to ABC transporter inhibition. Approximately 75% of dentate gyrus SP cells and 53% of SVZ SP cells were endothelial. This finding supports the work of Mouthon et al. (3) who showed that the side population of early postnatal SVZ cells contained cells positive for endothelial markers and did not possess NSC activity. The finding that hippocampal and subventricular endothelial cells formed the side population with effluxing capability is not surprising as, together with pericytes and astrocytes, they form and maintain the blood brain barrier (30–32). Accordingly, one of the main roles of endothelial cells is in brain homeostasis, which relies on the function of the ABC transporters (32). Primary adult cerebral endothelial cells show high ABC transporter protein levels (33), and single-cell RNA sequencing datasets from the human and mouse brain demonstrate that endothelial cells strongly express ABC transporter gene mRNA (34,35). Given that endothelial cells are often isolated using co-labeling of live cells with antibodies (24), our findings provide that using DCV is a less time-consuming, inexpensive, and an efficient alternative to isolate and study primary adult endothelial cells with active efflux.

### Requirements for optimization of side population assay

This study also showed that optimization of some parameters is required for accurate side population analysis. The optimization of the DCV dilution (Fig. S1) was done in order to avoid use of low or excess concentrations of DNA-binding dyes that can lead to the false identification of low-fluorescence cells as belonging to the side population (28), as non-effluxing cells in side and main populations are often continuous. Ensuring proper excitation with optimal voltage parameters for primary brain cells (S Fig. 1C, F) is also important to capture full heterogeneity of their DNA content. In addition, the usefulness of the relative cell size parameter (FSC) cannot be understated when locating very small side populations, such as those in the dentate and SVZ. The incorporation of the ABC transporter inhibitors further allows for more precise, higher resolution identification of the *bona fide* side population. This is demonstrated by the strong evidence of efflux within the side populations of dentate gyrus and SVZ cells, with only 0.34% and 0.22% cells not being inhibited by verapamil and fumitremorgin C, respectively, which may be due to the fact that our inhibitions do not include the ABCC family of transporters. Overall, our findings show that the dentate gyrus and SVZ side populations were modest in size and responded to verapamil and fumitremorgin C treatment but could only be identified while gating on DNA content together with relative cell size.

### Using SP assay to detect NSCs

Although the CD31+ endothelial cells represented most of the cells within the side population area of the dentate gyrus and SVZ, around 25% and 48% of cells in the respective side population did not express CD31. We hypothesize that these cells are not NSCs but may be small hematopoietic or microglial cells, as previously reported in the side populations of SVZ from early postnatal mice (3). Moreover, even if these cells are NSCs, given the majority of cells are endothelial cells, our data shows that the side population assay would not be efficient for the isolation of NSCs. This is in direct contrast to the efficiency of the side population assay to detect NSCs from cultured embryonic and adult cells from the SVZ niche (3,11). Such discrepancies point to biological differences in cultured and uncultured neurogenic cell niches, which has been suggested to be due to hypoxic conditions of neurosphere cultures (3,11). Independent of the cause of these differences, our findings and the work of others support that the identification of NSCs is limited to NSCs cultured *in vitro*. Additionally, we conclude that the use of DCV and analysis of the side population from *ex vivo* cell preparations from the neurogenic regions of the adult brain provides an inexpensive method to study endothelial cells.

## Acknowledgements

We would like to thank all members of the Lagace lab for continued valued feedback and funding provided by an NSERC discovery grant to DCL. We thank Vera A. Tang, the operations manager of the uOttawa Flow Cytometry and Virometry Core Facility for critical feedback and assistance with data collection.

## Supplementary Captions

**S Fig 1.**
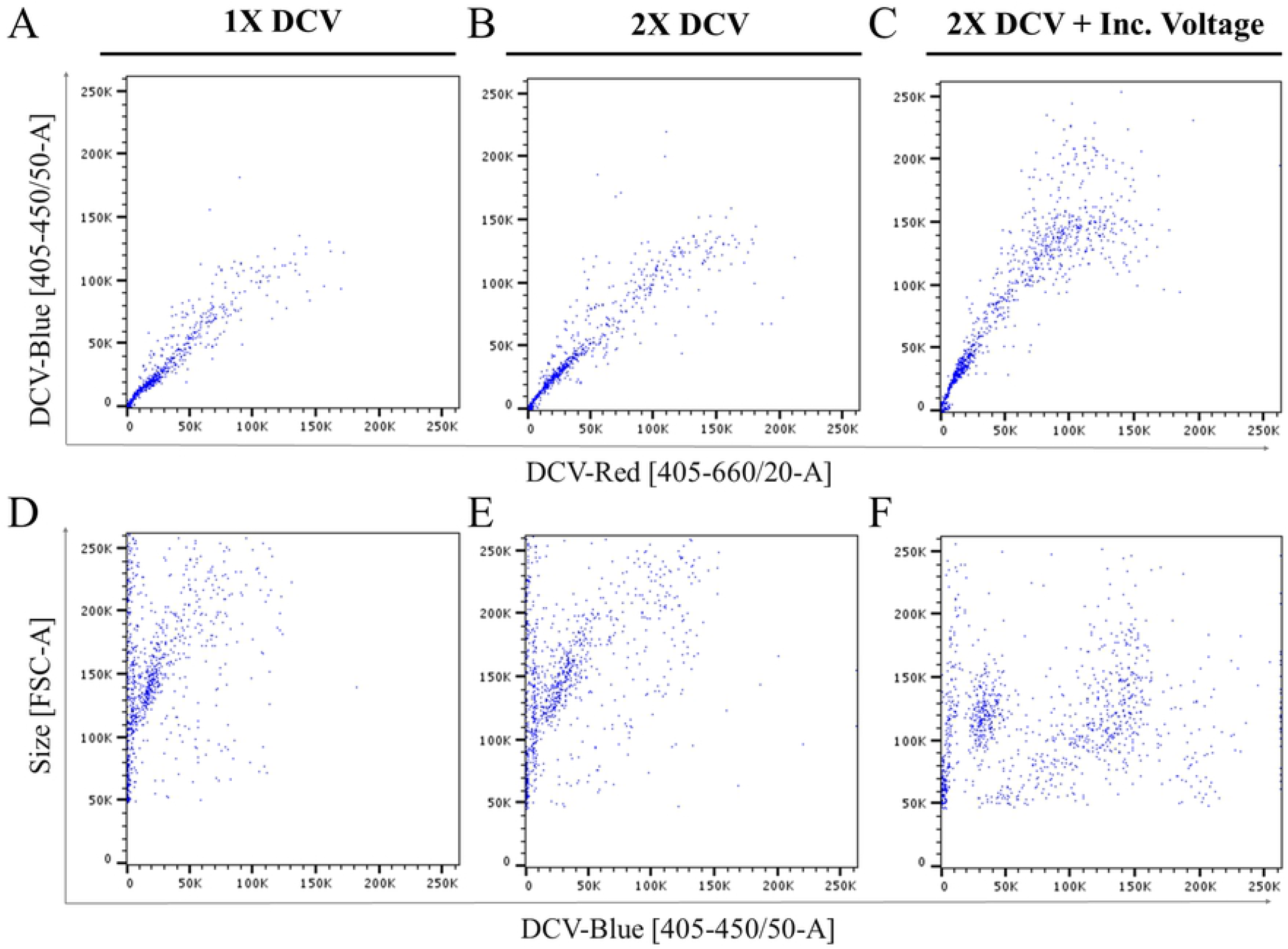
The titration of the DCV reagent in adult dentate gyrus cells. DCV staining testing 1X (A,D) or 2X DCV (B, E, C, F), as well as varying degrees of voltage for the 2X DCV concentration (B, E vs. C, F) as shown in dual fluorescence plots (A-C) and DCV-Blue/size plots (D-F). These results suggested 2X DCV with optimal excitation (C, F) was sufficient to distinguish the heterogeneous populations.

## References

1. Wu C-P, Zhou L, Xie M, Du H-D, Tian J, Sun S, et al. Identification of cancer stem-like side population cells in purified primary cultured human laryngeal squamous cell carcinoma epithelia. PLOS ONE. 2013 Jun 11;8(6):e65750.

2. Boesch M, Zeimet AG, Fiegl H, Wolf B, Huber J, Klocker H, et al. High prevalence of side population in human cancer cell lines. Oncoscience. 2016;3(3–4):85–7.

3. Mouthon M-A, Fouchet P, Mathieu C, Sii-Felice K, Etienne O, Lages CS, et al. Neural stem cells from mouse forebrain are contained in a population distinct from the “side population.” J Neurochem. 2006 Jan 1;99(3):807–17.

4. Salcido CD, Larochelle A, Taylor BJ, Dunbar CE, Varticovski L. Molecular characterisation of side population cells with cancer stem cell-like characteristics in small-cell lung cancer. Br J Cancer. 2010 May;102(11):1636–44.

5. Telford WG. Basic Cell Culture Protocols. Totowa: Humana Press; 2013.

6. Telford WG, Bradford J, Godfrey W, Robey RW, Bates SE. Side population analysis using a violet-excited cell-permeable DNA binding dye. Stem Cells. 2007;25(4):1029– 36.

7. Boesch M, Wolf D, Sopper S. Optimized stem cell detection using the DyeCycle-Triggered side population phenotype. Stem Cells International. 2015;2016:p e1652389.

8. Zhou S, Schuetz JD, Bunting KD, Colapietro A-M, Sampath J, Morris JJ, et al. The ABC transporter Bcrp1/ABCG2 is expressed in a wide variety of stem cells and is a molecular determinant of the side-population phenotype. Nat Med. 2001;7(9):1028–34.

9. Bond AM, Ming G, Song H. Ontogeny of adult neural stem cells in the mammalian brain. Curr Top Dev Biol; 2020;142:67–98.

10. Chaker Z, Codega P, Doetsch F. A mosaic world: puzzles revealed by adult neural stem cell heterogeneity. Wiley Interdiscip Rev Dev Biol. 2016;5(6):640–58.

11. Kim M, Morshead CM. Distinct populations of forebrain neural stem and progenitor cells can be isolated using side-population analysis. J Neurosci Off J Soc Neurosci. 2003 Nov 19;23(33):10703–9.

12. Pastrana E, Cheng L-C, Doetsch F. Simultaneous prospective purification of adult subventricular zone neural stem cells and their progeny. Proc Natl Acad Sci. 2009 Apr 14;106(15):6387–92.

13. Rietze RL, Valcanis H, Brooker GF, Thomas T, Voss AK, Bartlett PF. Purification of a pluripotent neural stem cell from the adult mouse brain. Nature. 2001;412(6848):736–9.

14. Murayama A, Matsuzaki Y, Kawaguchi A, Shimazaki T, Okano H. Flow cytometric analysis of neural stem cells in the developing and adult mouse brain. J Neurosci Res. 2002;69(6):837–47.

15. Hochgerner H, Zeisel A, Lönnerberg P, Linnarsson S. Conserved properties of dentate gyrus neurogenesis across postnatal development revealed by single-cell RNA sequencing. Nat Neurosci. 2018;21(2):290–9.

16. Shin J, Berg DA, Zhu Y, Shin JY, Song J, Bonaguidi MA, et al. Single-Cell RNA-Seq with Waterfall reveals molecular cascades underlying adult neurogenesis. Cell Stem Cell. 2015;17(3):360–72.

17. Dulken BW, Leeman DS, Boutet SC, Hebestreit K, Brunet A. Single-Cell transcriptomic analysis defines heterogeneity and transcriptional dynamics in the adult neural stem cell lineage. Cell Rep. 2017 Jan 17;18(3):777–90.

18. Artegiani B, Lyubimova A, Muraro M, van Es JH, van Oudenaarden A, Clevers H. A Single-Cell RNA sequencing study reveals cellular and molecular dynamics of the hippocampal neurogenic niche. Cell Rep. 2017;21(11):3271–84.

19. Hulspas R, Quesenberry PJ. Characterization of neurosphere cell phenotypes by flow cytometry. Cytometry. 2000;40(3):245–50.

20. Walker TL, Kempermann G. One mouse, two cultures: Isolation and culture of adult neural stem cells from the two neurogenic zones of individual mice. J Vis Exp. 2014 Feb 25;(84):e51225.

21. Hagihara H, Toyama K, Yamasaki N, Miyakawa T. Dissection of hippocampal dentate gyrus from adult mouse. J Vis Exp. 2009 Nov 17;(33): 1543.

22. Kannangara TS, Carter A, Xue Y, Dhaliwal JS, Béïque J-C, Lagace DC. Excitable adult-generated GABAergic neurons acquire functional innervation in the cortex after stroke. Stem Cell Rep. 2018 Dec 11;11(6):1327–36.

23. Babu H, Claasen J-H, Kannan S, Rünker AE, Palmer T, Kempermann G. A protocol for isolation and enriched monolayer cultivation of neural precursor cells from mouse dentate gyrus. Front Neurosci. 2011 Jul 14;5:89.

24. Crouch EE, Doetsch F. FACS isolation of endothelial cells and pericytes from mouse brain microregions. Nat Protoc. 2018;13(4):738–51.

25. Shaw TJ, Senterman MK, Dawson K, Crane CA, Vanderhyden BC. Characterization of intraperitoneal, orthotopic, and metastatic xenograft models of human ovarian cancer. Mol Ther. 2004 Dec;10(6):1032–42.

26. Tang Q-L, Liang Y, Xie X-B, Yin J-Q, Zou C-Y, Zhao Z-Q, et al. Enrichment of osteosarcoma stem cells by chemotherapy. Chin J Cancer. 2011;30(6):426–32.

27. Murase M, Kano M, Tsukahara T, Takahashi A, Torigoe T, Kawaguchi S, et al. Side population cells have the characteristics of cancer stem-like cells/cancer-initiating cells in bone sarcomas. Br J Cancer. 2009;101(8):1425–32.

28. Golebiewska A, Brons NHC, Bjerkvig R, Niclou SP. Critical appraisal of the side population assay in stem cell and cancer stem cell research. Cell Stem Cell. 2011 Feb 4;8(2):136–47.

29. Boesch M, Zeimet AG, Reimer D, Schmidt S, Gastl G, Parson W, et al. The side population of ovarian cancer cells defines a heterogeneous compartment exhibiting stem cell characteristics. Oncotarget. 2014 Jun 1;5(16):7027–39.

30. Giannoni P, Badaut J, Dargazanli C, De Maudave AF, Klement W, Costalat V, et al. The pericyte–glia interface at the blood–brain barrier. Clin Sci. 2018 Feb 14;132(3):361–74.

31. Ayloo S, Gu C. Transcytosis at the blood–brain barrier. Curr Opin Neurobiol. 2019 Aug 1;57:32–8.

32. Mahringer A, Fricker G. ABC transporters at the blood–brain barrier. Expert Opin Drug Metab Toxicol. 2016 May 3;12(5):499–508.

33. Warren MS, Zerangue N, Woodford K, Roberts LM, Tate EH, Feng B, et al. Comparative gene expression profiles of ABC transporters in brain microvessel endothelial cells and brain in five species including human. Pharmacol Res. 2009 Jun 1;59(6):404–13.

34. Song HW, Foreman KL, Gastfriend BD, Kuo JS, Palecek SP, Shusta EV. Transcriptomic comparison of human and mouse brain microvessels. Sci Rep. 2020 Jul 23;10(1):12358.

35. Kiss T, Nyúl-Tóth Á, Balasubramanian P, Tarantini S, Ahire C, DelFavero J, et al. Single-cell RNA sequencing identifies senescent cerebromicrovascular endothelial cells in the aged mouse brain. GeroScience. 2020 Mar 31;42(2):429–44.

